# Circadian rhythms have significant effects on leaf-to-canopy gas exchange under field conditions

**DOI:** 10.1101/054593

**Authors:** Víctor Resco de Dios, Arthur Gessler, Juan Pedro Ferrio, Josu G Alday, Michael Bahn, Jorge del Castillo, Sébastien Devidal, Sonia García-Muñoz, Zachary Kayler, Damien Landais, Paula Martín, Alexandru Milcu, Clément Piel, Karin Pirhofer-Walzl, Olivier Ravel, Serajis Salekin, David T Tissue, Mark G Tjoelker, Jordi Voltas, Jacques Roy

**Affiliations:** Department of Crop and Forest Sciences-AGROTECNIO Center, Universitat de Lleida, 25198 Lleida, Spain.; Hawkesbury Institute for the Environment, University of Western Sydney, Richmond, NSW 2753, Australia.; Swiss Federal Institute for Forest, Snow and Landscape Research WSL Long-term Forest Ecosystem Research (LWF), 8903 Birmensdorf, Switzerland.; Institute for Landscape Biogeochemistry, Leibniz-Centre for Agricultural Landscape Research (ZALF), 15374 Müncheberg, Germany.; School of Environmental Sciences, University of Liverpool, Liverpool, L69 3GP, UK.; Institute of Ecology, University of Innsbruck, 6020 Innsbruck, Austria.; Ecotron Européen de Montpellier, UPS 3248, CNRS, Campus Baillarguet, 34980, Montferrier-sur-Lez, France.; IMIDRA, Finca “El Encín”, 28800 Alcalá de Henares, Madrid, Spain.; CNRS, Centre d’Ecologie Fonctionnelle et Evolutive (CEFE UMR 5175), 1919 route de Mende, F-34293 Montpellier, France; Erasmus Mundus Master on Mediterranean Forestry and Natural Resources Management, Universitat de Lleida, 25198 Lleida, Spain.

**Keywords:** circadian clock, ecological memory, net ecosystem exchange, scaling, stomatal conductance and models, photosynthesis, transpiration

## Abstract

Molecular clocks drive oscillations in leaf photosynthesis, stomatal conductance and other cell and leaf level processes over ~24 h under controlled laboratory conditions. The influence of such circadian regulation over whole canopy fluxes remains uncertain and diurnal CO_2_ and H_2_O vapor flux dynamics in the field are currently interpreted as resulting almost exclusively from direct physiological responses to variations in light, temperature and other environmental factors. We tested whether circadian regulation would affect plant and canopy gas exchange at the CNRS Ecotron. Canopy and leaf level fluxes were constantly monitored under field-like environmental conditions, and also under constant environmental conditions (no variation in temperature, radiation or other environmental cues). Here we show first direct experimental evidence at canopy scales of circadian gas exchange regulation: 20-79% of the daily variation range in CO_2_ and H_2_O fluxes occurred under circadian entrainment in canopies of an annual herb (bean) and of a perennial shrub (cotton). We also observed that considering circadian regulation improved performance in commonly used stomatal conductance models. Overall, our results show that overlooked circadian controls affect diurnal patterns of CO_2_ and H_2_O fluxes in entire canopies and in field-like conditions, although this process is currently unaccounted for in models.

## Introduction

Terrestrial ecosystems play a major role in the global carbon and water cycles. It is currently estimated that ~30% of fossil fuel emissions are sequestered by land (Canadell *et al.*, 2007), and that ~60% of annual precipitation is returned to the atmosphere through evapotranspiration, a flux largely dominated by transpiration (Schlesinger and Jasechko, 2014). There is a long tradition of research within the Earth Sciences on deciphering the mechanisms underlying diurnal variations in photosynthesis and transpiration (Jones, 1998; Chapin *et al.*, 2002; Sellers *et al.*, 1997; Hollinger *et al.*, 1994). This research has mostly focused on direct physiological responses to the environment. That is, towards understanding how the photosynthetic machinery and stomatal function respond and react to changes in radiation, temperature, vapor pressure deficit, and other environmental drivers.

A significantly smaller body of research has sought to disentangle whether, apart from responses to exogenous factors, endogenous processes could also play a role (Resco *et al.*, 2009). It has been documented, for instance, how for a given level of water potential and concentration of abscisic acid (ABA), stomatal conductance is higher in the morning than in the afternoon (Mencuccini *et al.*, 2000). The process controlling this phenomenon is the circadian clock (Mencuccini *et al.*, 2000), an endogenous timer of plant metabolism that controls the temporal pattern of transcription in photosynthesis, stomatal opening, and other physiological processes (Hubbard and Webb, 2015). There are additional processes creating endogenous flux variation, but only the circadian clock will be addressed here.

Research on the regulation of photosynthesis and transpiration within field settings by the circadian clock is much smaller than research on direct responses to the environment. For instance, we conducted a literature search on the database Scopus (3^rd^ March 2016) with the words “circadian AND ecosystem AND photosynthesis” in the title or abstract and we obtained 11 results. This is contrast with the 3,367 results found with the words “ecosystem AND photosynthesis”, or with the 1,085 results with the words “temperature AND ecosystem AND photosynthesis”. The few studies that do mention circadian regulation, often consider it as a negligible driver at canopy or ecosystem scales (Lasslop *et al.*, 2010; Williams *et al.*, 2014), although there are a few notable exceptions (Dietze, 2014; Stoy *et al.*, 2014).

The explicit statement that circadian regulation is a negligible driver of gas exchange in the field has its roots in a study conducted almost twenty years ago and entitled “Circadian rhythms have insignificant effects on plant gas exchange under field conditions” (Williams and Gorton, 1998). This was a pioneer study that, for the first time, took research on circadian rhythms outside of lab settings and worked with a non-model species from wetland and understory environments (*Saururus cernuus* L.). The elegant study from Williams and Gorton (1998) measured leaf level fluxes under “constant environmental conditions” (that is, when temperature, radiation and other environmental drivers do not change through time). They documented a 24-h oscillation in gas exchange within growth chambers, consistent with circadian regulation of gas exchange. They then tested whether such circadian regulation would be also significant in the field by adding a sinusoidal variation to a biochemical model of gas exchange. Under these conditions, they observed how model goodness-of-fit increased, but only by 1%. Hence they concluded that circadian regulation of gas exchange in the field was insignificant. That study was focused on photosynthesis and, as we write, we are not aware of any attempts to include circadian regulation into stomatal conductance models.

Besides Williams and Gorton (1998), others have attempted to infer circadian regulation of gas exchange in the field by filtering flux tower data and obtained circumstantial evidence that circadian regulation could indeed be an important driver of net ecosystem exchange in the field (Resco de Dios *et al.*, 2012; Doughty *et al.*, 2006), and also of isoprene emissions (Hewitt *et al.*, 2011). Others, working with nocturnal transpiration, have additionally documented how circadian regulation over nocturnal stomatal conductance affects the transpiration stream in whole-trees (Resco de Dios *et al.*, 2013) or even entire plant canopies (Resco de Dios *et al.*, 2015).

However, direct tests of circadian regulation of photosynthesis and of daytime transpiration at canopy scales are still missing. Understanding whether or not circadian regulation in gas exchange scales up into canopies is important to understand the potential implications of the circadian clock as a driver of diurnal flux dynamics, and there are reasons to expect a dilution of circadian effects as we move up in scale. In mammals, a hierarchical network of circadian clocks exists, with a unique central oscillator on the suprachiasmatic nucleus in the brain (Endo, 2016). However, circadian clocks in plants are more autonomous and there is little evidence that the clock in different leaves is synchronized (Endo, 2016). Circadian rhythms are entrained by environmental cues of light and temperature. Therefore, at canopy scale, different leaves will experience different light and temperature cues and we could observe uncoupled circadian rhythms in different leaves within and across plants, potentially diluting any circadian effects at canopy scales.

These are the research gaps addressed by this study. We monitored leaf and canopy gas exchange under field-like and also under constant environmental conditions in bean (*Phaseolus vulgaris)* and cotton (*Gossypium hirsutum*) canopies within an experimental Ecotron and tested: i) whether circadian regulation in photosynthesis and daytime stomatal conductance scales up from leaves to canopy; and ii) whether adding a circadian oscillator into well-known stomatal models would significantly increase model fit.

## MATERIALS AND METHODS

### Ecotron and general experimental set-up

The experiment was performed at the Macrocosms platform of the Montpellier European Ecotron, Centre National de la Recherche Scientifique (CNRS, France). We used 12 outdoor macrocosms (6 planted with bean and 6 with cotton) where the main abiotic (air temperature, humidity and CO_2_ concentration) drivers were automatically controlled. In each macrocosm, plants were grown on a soil (area of 2 m^2^, depth of 2 m) contained in a lysimeter resting on a weighing platform. The soil was collected from the flood plain of the Saale River near Jena, Germany, and used in a previous Ecotron experiment on biodiversity (Milcu *et al.*, 2014). After that experiment, the soil was ploughed down to 40 cm and fertilized with 25/25/35 NPK (MgO, SO_3_ and other oligoelements were associated in this fertilizer: Engrais bleu universel, BINOR, Fleury-les-Aubrais, FR).

The soil was regularly watered to *ca*. field capacity by drip irrigation, although irrigation was stopped during each measurement campaign (few days) to avoid interference with water flux measurements. However, no significant differences (at *P* < 0.05, paired t-test, n=3) in leaf water potential occurred between the beginning and end of these measurement campaigns, indicating no effect of a potentially declining soil moisture on leaf hydration (Resco de Dios *et al.*, 2015).

Environmental conditions within the macrocosms (excluding the experimental periods) were set to mimic outdoor conditions, but did include a 10% light reduction by the macrocosm dome cover. During experimental periods, light was controlled by placing a completely opaque fitted cover on each dome to block external light inputs (PVC coated polyester sheet Ferrari 502, assembled by IASO, Lleida, Spain), and by using a set of 5 dimmable plasma lamps (GAN 300 LEP with the Luxim STA 41.02 bulb, with a sun-like light spectrum, Fig. S1); these lamps were hung 30 cm above the plant canopy and provided a PAR of 500 µmol m^−2^ s^−1^ (the maximum possible by those lamps). We measured PAR at canopy level with a quantum sensor (Li-190, LI-COR Biosciences, Lincoln, NE, USA) in each macrocosm.

**Figure. S1:**
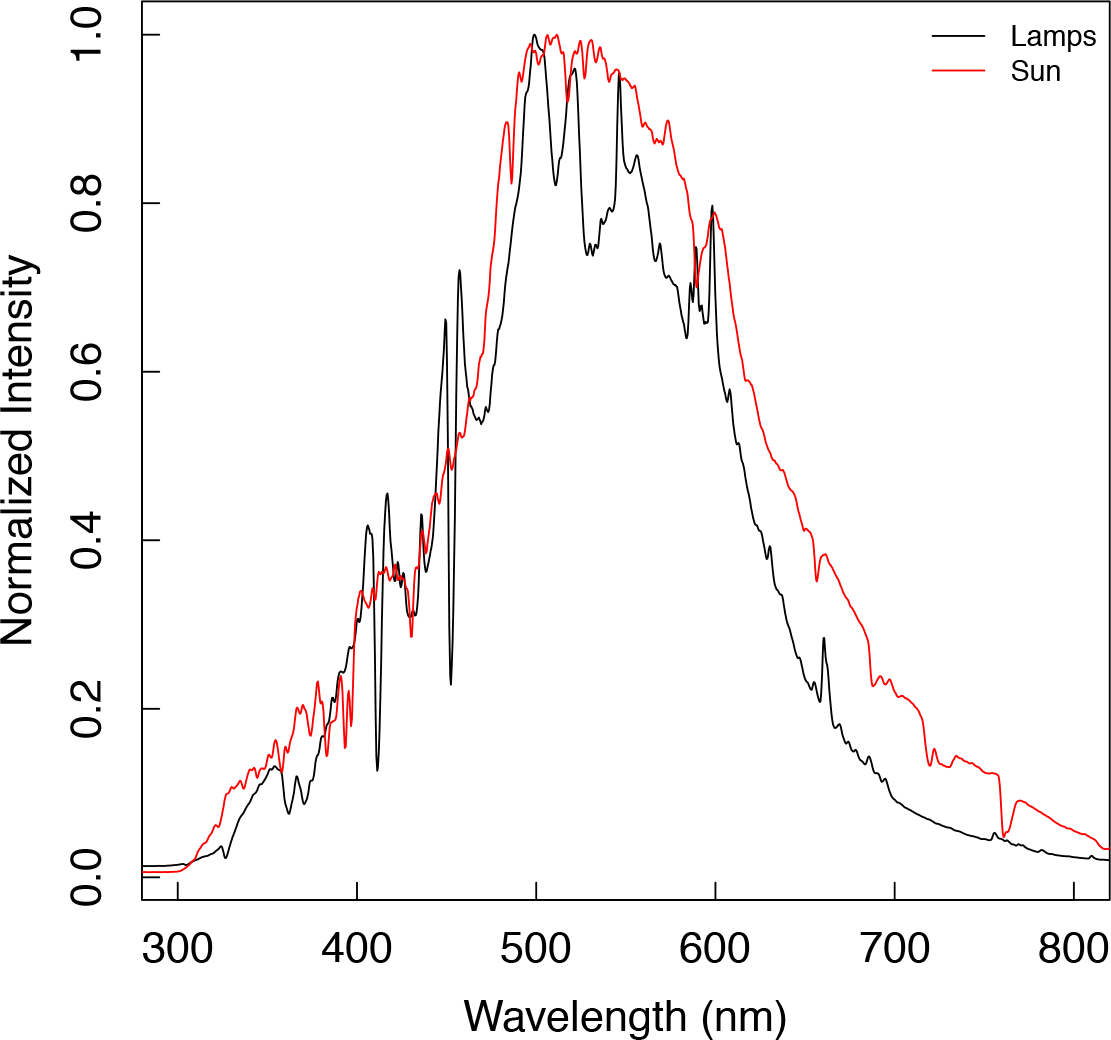
The plasma lamps used in the experiment had a sun-light spectrum. Intensity at each wavelength was measured with a Jaz spectrometer (Ocean Optics UV-NIR detector, Jasper, GA, USA).

Bean and cotton were planted in 5 different rows within the domes on 10^th^ July 2013, one month before the start of the measurements, and thinned to densities of 10.5 and 9 individuals m^−2^, respectively. Cotton (STAM-A16 variety by INRAB/CIRAD) is a perennial shrub with an indeterminate growth habit. This cotton variety grows to 1.5-2 m tall and has a pyramidal shape and short branches. Bean (recombinant inbred line RIL-115 bred by INRA Eco&Sol) is an annual herbaceous species. RIL-115 is a fast growing, indeterminate dwarf variety, 0.3-0.5 m tall; it was inoculated with *Rhizobium tropici* CIAT 899 also provided by INRA.

### Measuring techniques

Each unit of the Macrocosms platform was designed as an open gas exchange system to continuously measure CO_2_ net ecosystem exchange by measuring the air flow at the inlet of each dome (thermal mass flowmeter Sensyflow iG, ABB, Zurich, CH) and by sequentially (every 12 min) measuring the CO_2_ concentration at each inlet and outlet using a multiplexer system coupled with two LI-7000 CO_2_/H_2_O analyzers (LI-COR Biosciences, Lincoln, NE, USA). Belowground fluxes were prevented from mixing with canopy air by covering the soil with a plastic sheet during the entire experimental period. Substantial internal air mixing within the dome (2 volumes per min) reduced the canopy boundary layer and minimized the CO_2_ concentration gradients within the dome. A slight atmospheric over-pressure (5 to 10 Pa) applied to the plastic sheet (through the slits made for the plant stems) covering the soil minimized potential mixing of soil respiration fluxes with aboveground fluxes. Indeed, we observed negligible CO_2_ flux at the onset of the experiment (immediately after seed germination, when there was no significant carbon assimilation), indicating lack of significant CO_2_ efflux on the canopy above the plastic sheet. Transpiration was measured continuously by weighing lysimeters with four shear beam load cells per lysimeter (CMI-C3 Precia-Molen, Privas, France), and calculated from the slope of the temporal changes in mass using a generalized additive model with automated smoothness selection (Wood, 2006).

For each crop, three macrocosms were dedicated to leaf level measurements (researchers entered periodically) and the remaining three macrocosms were ‘undisturbed’ and dedicated to continuous canopy gas exchange measurements. During the experiment, bean and cotton generally remained at the inflorescence emergence developmental growth stage (Munger *et al.*, 1998; codes 51-59 in BBCH scale, the standard phenological scale within the crop industry; Feller *et al.*, 1995). Further details on Ecotron measurements have been provided elsewhere (Resco de Dios *et al.*, 2015; Milcu *et al.*, 2014).

We measured leaf gas exchange using a portable photosynthesis system (LI-6400XT, Li-Cor, Lincoln, Nebraska, USA), after setting the leaf cuvette to the same environmental conditions as the macrocosms. We conducted spot gas exchange measurements every 4 hours in three leaves within each macrocosm, and average values for each of the 3 macrocosms per species were used in subsequent analyses. Different leaves from different individuals were measured during each measurement round. Leaf temperature was independently measured at the time of gas exchange measurements with an infra-red thermometer (MS LT, Optris GmbH, Berlin, Germany) and no significant difference with air temperature recorded by the *T*_air_ probe (PC33, Mitchell Instrument SAS, Lyon, France) was observed (intercept = −4.3 ± 4.5 [mean ± 95%CI]; slope = 1.15 ± 0.17; R^2^ = 0.89).

#### Question 1: Does circadian regulation scale up to affect whole canopy fluxes?

We tested whether leaf circadian regulation scaled up to affect whole ecosystem CO_2_ and H_2_O fluxes by examining leaf carbon assimilation (*A*_1_) and stomatal conductance (*g*_s_), in addition to canopy carbon assimilation (*A*_c_) and transpiration (*E*_c_) under “constant” and “changing” environmental conditions. Canopies were originally entrained (“changing” conditions) by mimicking the temporal patterns in *T*_air_ (28/19 °C, max/min) and VPD (0.5/1.7 kPa) of an average sunny day in August in Montpellier (Fig. 1). Photoperiod was set to 12 h of darkness and 12 h of light during entrainment, and a maximum PAR of 500 μlmol m^−2^ s^−1^ at canopy height was provided by the plasma lamps. This radiation is substantially lower than in a sunny day in Montpellier, but we do not know of any facility in the world that allows for environmental control and automated flux measurements at canopy scales under a higher radiation due to technical limitations. After a 5-day entrainment period, we maintained PAR, *T*_air_ and VPD constant for 48-h starting at solar noon (“constant” conditions). These experiments were performed between 8^th^ August and 3^rd^ September 2013.

**Fig. 1.**
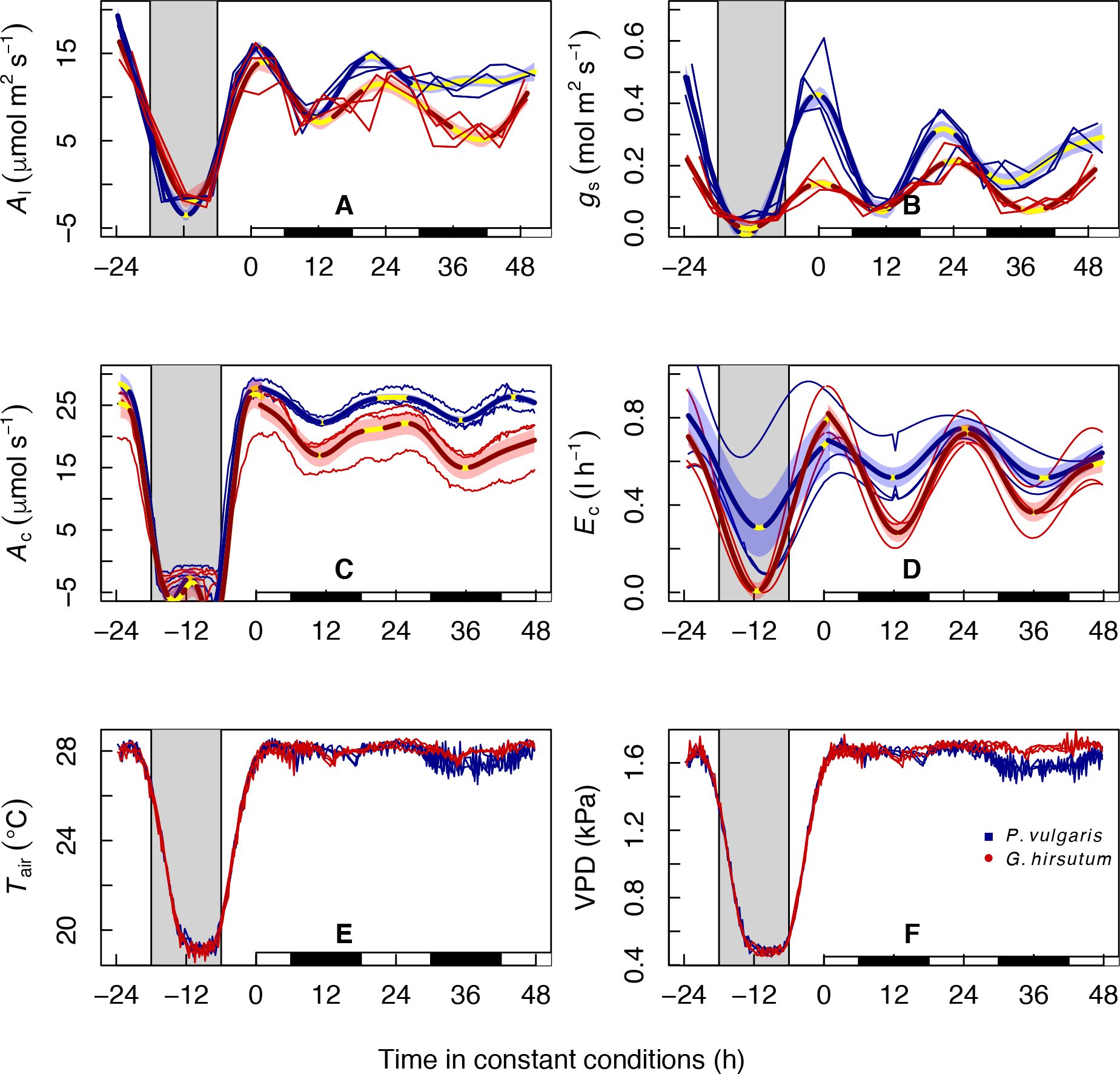
Circadian regulation of leaf and canopy-scale fluxes of CO_2_ and H_2_O. Environmental conditions of Temperature (*T*_air_) and Vapor Pressure Deficit (VPD) mimicked an average August day in Montpellier, with 500 µmol m^−2^ s^−1^ PAR (first 24 h shown), and remained constant for the following 48 h starting at solar noon. The grey (white) background indicates when PAR was at (above) 0 µmol m^−2^ s^−1^. The white and black rectangles at the base indicate the subjective day (when it would have been daytime during entrainment) and subjective night, respectively, under constant conditions. Thin lines represent measured values at each of three replicate macrocosms, and thick lines (and shaded error intervals) indicate the prediction (and SE) of Generalized Additive Mixed Model (GAMM) fitting separately for each species (some lines may overlap). Significant variation (GAMM best-fit line portions not yellow) in leaf and canopy carbon assimilation (*A*_l_ and *A*_c_, respectively), in stomatal conductance (*g*_s_) and canopy transpiration (*E*_c_), as well as in their ratios prevailed for all fluxes and processes at least in the first 24 h under constant conditions. This can be fully attributed to circadian action. Clock regulation is plastic and may relax after prolonged exposures to constant conditions (Hennessey *et al.*, 1993). Negative dark-time values of *A*_l_/*g*_s_ and *A*_c_*/E*_c_ were cropped as they lack biological meaning.

We examined statistical significance of temporal patterns with Generalized Additive Mixed Model (GAMM) fitted with automated smoothness selection (Wood, 2006) in the R software environment (*mgcv* library in R 3.1.2, The R Foundation for Statistical Computing, Vienna, Austria), including macrocosms as a random factor. This approach was chosen because it makes no *a priori* assumption about the functional relationship between variables. We accounted for temporal autocorrelation in the residuals by adding a first-order autoregressive process structure (*nlme* library (Pinheiro and Bates, 2000)). Significant temporal variation in the GAMM best-fit line was analyzed after computation of the first derivative (the slope, or rate of change) with the finite differences method. We also computed standard errors and a 95% point-wise confidence interval for the first derivative. The trend was subsequently deemed significant when the derivative confidence interval was bounded away from zero at the 95% level (for full details on this method see Curtis and Simpson, 2014). Non-significant periods, reflecting lack of local statistically significant trending, are illustrated on the figures by the yellow line portions, and significant differences occur elsewhere. The magnitude of the range in variation driven by the circadian clock (Table 1) was calculated using GAMM maximum and minimum predicted values.

**Table 1:** Quantification of the circadian-driven range in variation of diurnal gas exchange. The variation in fluxes attributable to the clock in Fig. 1 was derived from the ratio between the range (maximum GAMM predicted value minus minimum GAMM predicted value) in each flux while keeping environmental conditions constant (last 48 h in Fig. 1), divided by the range during the entrainment phase (first 24 h in Fig. 1). Although nocturnal stomatal conductance and transpiration were always above 0 during entrainment, even during dark periods, we forced their minimum to be zero for this calculation. This increased the magnitude of the variation during entrainment, thus leading to under-estimations of the % variation attributable to the clock. Nocturnal carbon assimilation was also fixed at 0, because no C assimilation occurs in the dark.

| Process | Species | Scale | Variation during entrainment |  |  | Variation during constant conditions |  |  | % clock-driven variation |
| --- | --- | --- | --- | --- | --- | --- | --- | --- | --- |
|  |  |  | Max (SE) | Min | Max-Min | Max (SE) | Min (SE) | Max-Min |  |
| Carbon assimilation | *P.* | Leaf ($\mu\text{molm}^{-2}\text{s}^{-1}$) | 19.30 (0.97) | 0 | 19.30 | 15.67 (0.66) | 7.79 (0.63) | 7.88 | 40.83 |
| | *vulgaris* | Ecosystem ($\mu\text{molm}^{-2}\text{s}^{-1}$) | 28.42 (1.74) | 0 | 28.42 | 27.84 (0.64) | 22.25 (0.61) | 5.59 | 19.67 |
| | *G.* | Leaf ($\mu\text{molm}^{-2}\text{s}^{-1}$) | 16.32 (1.42) | 0 | 16.32 | 14 (0.80) | 5.13 (0.84) | 8.87 | 54.35 |
| | *hirsutum* | Ecosystem ($\mu\text{molm}^{-2}\text{s}^{-1}$) | 26.76 (2.23) | 0 | 26.76 | 25.03 (1.82) | 14.96 (1.81) | 10.07 | 37.63 |
| Water fluxes | *P.* | Leaf (conductance, $\text{molm}^{-2}\text{s}^{-1}$) | 0.48 (0.04) | 0 | 0.48 | 0.43 (0.03) | 0.05 (0.03) | 0.38 | 79.17 |
| | *vulgaris* | Ecosystem ($\text{l h}^{-1}$) | 0.81 (0.13) | 0 | 0.81 | 0.75 (0.07) | 0.52 (0.07) | 0.23 | 28.39 |
| | *G.* | Leaf (conductance, $\text{molm}^{-2}\text{s}^{-1}$) | 0.22 (0.02) | 0 | 0.22 | 0.21 (0.01) | 0.05 (0.01) | 0.16 | 72.73 |
| | *hirsutum* | Ecosystem ($\text{l h}^{-1}$) | 0.79 (0.08) | 0 | 0.79 | 0.79 (0.06) | 0.28 (0.06) | 0.51 | 64.55 |

#### Question 2: Does adding a circadian oscillator improve the performance of stomatal models?

The stomatal models were fitted with non-linear least squares regression using the base R packages. The models used (Medlyn *et al.*, 2011; Leuning, 1995; Ball *et al.*, 1987) have two common fitting parameters, which we will call *g*_0_ (minimal conductance, or the intercept of the model) and *g*_1_ (slope, that relates *g*_s_ to *A*_1_ and environmental variables). We ran the models with and without *g*_0_, as the interpretation of minimal conductance remains elusive (Medlyn *et al.*, 2011). We observed changing *A*_1_/*g*_s_, so circadian oscillations were added to modify the values of *g*_1_ over time:

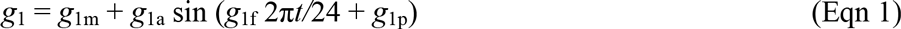

where subscripts *m*, *a*, *f* and *p* indicate the mean *g*_1_ value, the amplitude, frequency and phase of the rhythm, respectively, and *t* is time in hours (since experiment onset). That is, we studied the clock effect on *g*_s_ model predictions by comparing the original model formulations (Medlyn *et al.*, 2011; Leuning, 1995; Ball *et al.*, 1987) before (without circadian oscillator), and after (with circadian oscillators) replacing *g*1 in the original formulations by Eqn 1. We derived *g*_1m_ for models that included a circadian oscillator from the estimate of *g*_1_ in the corresponding models without a circadian oscillator, and the frequency (*g*_1f_) was additionally fixed at 24 h (*g*_1f_ = 1).

We conducted three different model runs for each of the three different models of stomatal conductance. First, each *g*_s_ model was calibrated and validated with the entire leaf-level dataset (Fig. 1). Second, we calibrated each model under changing diurnal conditions of PAR, *T*_air_ and VPD (first 24 h in Fig. 1) and validated it with data under constant PAR, *T*_air_ and VPD conditions (last 48 h in Fig. 1). Third, we calibrated each model under constant PAR, *T*_air_ and VPD conditions, and validated it with data under changing PAR, *T*_air_ and VPD. Given the distinctly different pattern of environmental conditions during the changing and constant phases, the last two model runs were included to represent changes in model fit under ‘novel’ environmental conditions. Importantly, the third model run would be comparable with the study of Williams and Gorton (1998), in that it would it use data under constant environmental conditions to infer the effect over changing environmental conditions.

The models were fitted independently for each species, but observed and predicted values were then combined for validation. We calculated R^2^ from the regression between observed vs predicted values, and Akaike Information Criterion (AIC) was obtained as:

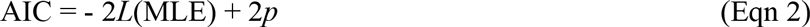

where *L*(MLE) is the likelihood function evaluated at the maximum likelihood estimates, and *p* the number of parameters. AIC reduction (ΔAIC) for a model was calculated from the difference to the smallest AIC, and the weights (*w*_i_) from the ratio between the relative likelihood of a model 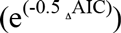 to the sum of all relative likelihoods.

## RESULTS

### Circadian regulation scales up to affect whole canopy fluxes

We entrained the bean and cotton canopies for 5 days under average daily patterns of air temperature (*T*_air_) and vapour pressure deficit (VPD) for an August day in Montpellier, albeit with lower photosynthetically active radiation (PAR, up to 500 µmol m^−2^ s^−1^, Fig. 1 E-F). Thereafter, we kept *T*_air_, VPD and PAR constant for 48 h and throughout this time-course, we observed continuous temporal variation in leaf-level and integrated canopy carbon dioxide (*A*) and water vapour (*E*) fluxes between 20-79% of the range observed during entrainment (depending on flux, species and scale, full details in Fig. 1, Table 1). Temporal variations of *A* and *E* at the leaf- and canopy-levels under a constant environment showed a period of ~24 h, consistent with circadian regulation of leaf photosynthesis (*A*_l_) and stomatal conductance (*g*_s_).

There were some subtle differences across species in terms of the magnitude of the oscillation but, overall, similar patterns were observed. There was a slight dilution of circadian regulation as we moved up in scale. For instance, the magnitude of the clock driven variation was 41-54% for *A*_l_, but 20-38% in *A*_c_. Similary, while *g*_s_varied by 72-79% under constant conditions, the variation in *E*_c_ was 28-64%. However, despite this dilution, we always observed a significant self-sustained 24-h oscillation in *A*_c_ as well as in *E*_c_.

It could be argued that this calculation of the importance of circadian regulation will tend to overestimate its importance because it is based upon a 24 h cycle whereas in reality no *A*_c_ occurs during the night, and *E*_c_ will be lower under a normal night (when it is dark) than in the subjective night in the free running period. We thus re-calculated the magnitude of the oscillation in *A*_c_ and *E*_c_ only during the 12 h of the subjective day in the free running period and observed that it was 15.4% and 24.0%, respectively, for bean, and 29.75 and 37.7%, respectively, for cotton.

### A circadian oscillator improves the performance of stomatal models

As previously mentioned, we conducted different model runs by varying the calibration and validation datasets. Depending on the combination of the datasets, we observed that either variations from the models originally proposed either by Medlyn *et al.* (2011) or by Leuning (1995) performed the best (Table 2). However, regardless of the dataset, the best model was always one that included a circadian oscillator in the slope (Table 2). This result indicates that inclusion of a circadian oscillator significantly improves model performance. This was also the case when using the conditions closer to the experiment by Williams and Gorton (1998)

**Table 2:**
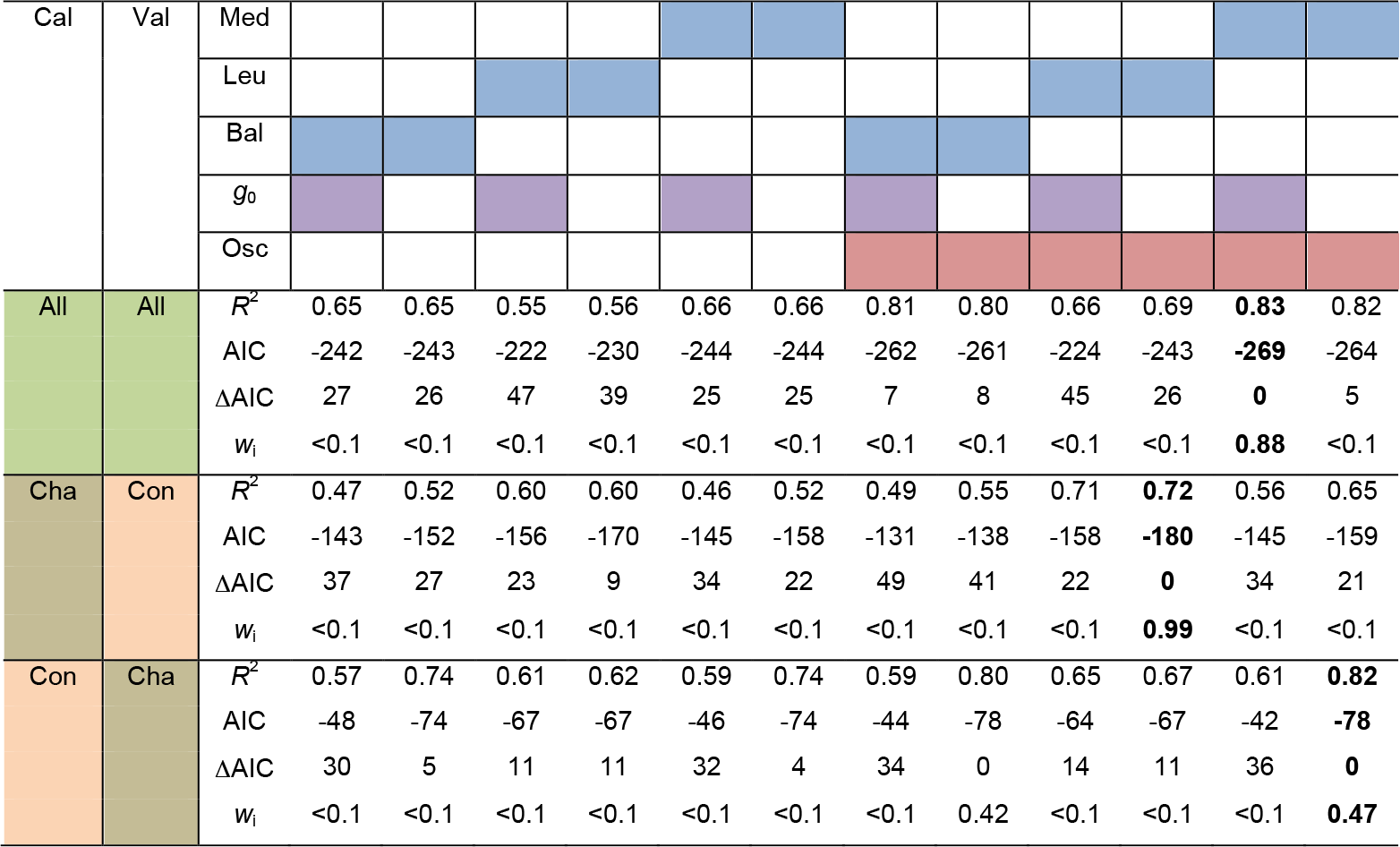
Model fits of leaf stomatal conductance improve with inclusion of a circadian oscillator. Results of fitting the models of stomatal conductance proposed by Medlyn *et al.* 2011 (Med), Leuning 1995 (Leu), and Ball *et al.* 1987 (Bal, indicated in blue), excluding and including minimal conductance (*g*_0_, in purple, a fitting parameter across models, see Methods), and excluding and including a circadian oscillator (Osc, in red). Data used for calibration (Cal) and validation (Val) are indicated by the colors green (entire dataset from Fig. 1B, All), brown (under changing conditions in Fig. 1B, Cha), or orange (under constant conditions in Fig. 1B, Con). Values in bold indicate the best-fit model for each combination of calibration/validation datasets. Models were assessed by their *R*^2^, the Akaike Information Criterion (AIC), AIC reduction (ΔAIC) and the weight of each model (*w*_i_). The model with the smallest ΔAIC and largest *w*_i_ is considered the most plausible (Burnham and Anderson, 2002). Regardless of the dataset, inclusion of a circadian oscillator rendered the models more plausible.

## DISCUSSION

We observed how, in the absence of fluctuations in environmental drivers, *A* and *E* both oscillated significantly. There is a myriad of endogenous processes that could affect temporally carbon and water fluxes, such as carbohydrate accumulation (Azcón-Bieto, 1983) or hydraulic feedbacks (Jones, 1998), to name a couple. However, these feedbacks will generally tend towards decreasing *A* and *E* over time. The only mechanism currently known to create a self-sustained 24h cycle is the circadian clock (McClung, 2006; Müller *et al.*, 2014).

It is well-known that radiation is the major environmental driver of as exchange, and it could create 100% of the diurnal oscillation. *T*_air_ and VPD are often considered as the next most important environmental drivers of diurnal flux dynamics. Although we measured neither *T*_air_ nor VPD responses alone during these experiments, other studies with these species typically document that, in the absence of strong environmental stress, *T*_air_ and VPD responses could lead to diurnal flux variation of the same order of magnitude as those observed in this study(Duursma *et al.*, 2014). In other words, the oscillation in *A*_c_ and *E*_c_ observed in this study (Table 1) would be comparable to that documented in *T*_air_ or VPD response curves.

To more fully understand the up-scaling of circadian rhythms, we need to explore further how canopy structure affects ecosystem-level expression of circadian regulation. Circadian regulation in understory species has been shown to be less important than in overstory species (Doughty *et al.*, 2006), presumably because the predictability of environmental cues diminishes under a canopy. An ecosystem-level analogy would be forests with high leaf area index, where a relatively large proportion of carbon fixation and water loss may be conducted by shaded leaves. In fact, we always observed a higher degree of circadian-driven variation in leaf level compared to canopy level fluxes (Table 1), which could have resulted from the larger proportion of shaded leaves at the canopy scale. Greater understanding of the relative importance of circadian regulation on ecosystem processes, as a function of leaf canopy structure, should thus be a future research objective.

We also conducted a modeling exercise where *g*_s_ was calibrated with the constant conditions dataset and then validated under changing conditions, which would be similar to the approach by Williams and Gorton (1998). However, although validation did not occur under strictly field conditions, it did occur under field-like conditions. Since we observed significant improvements in mode fits, we can conclude that the assertion of circadian rhythms having insignificant rhythms for gas exchange under field settings needs to be revised.

Circadian regulation had a more important effect on stomatal conductance and ecosystem transpiration than on leaf and canopy carbon assimilation (Fig. 1). This is probably the reason why circadian regulation here significantly improved stomatal model output here, while this was may not have been the case in previous studies on photosynthesis (Williams and Gorton, 1998). It is worth noting that there are many reports of a hysteresis on tree transpiration such that, for a given environmental condition, transpiration is higher in the morning than in the afternoon (Zhang *et al.*, 2014; Tuzet *et al.*, 2003; O'Grady *et al.*, 1999). This phenomenon has been often explained in terms of hydraulic feedbacks on stomata. However, our results, along with further experiments on circadian regulation of stomata (Mencuccini *et al.*, 2000; Marenco *et al.*, 2006), indicate that circadian rhythmicity could be another factor that, at least partly, explains hysteretic water fluxes.

Here, we have used an empirical approach that considers time as a surrogate of circadian regulation. Importantly, we observed how the circadian oscillator enhanced the performance of diurnal leaf-level stomatal models (Table 2). However, we acknowledge that the use of time as a surrogate for circadian action is not fully satisfactory; yet, at present, this is the only approach given limited understanding of circadian processes at the scale of relevance for this analysis.

Previous studies have shown that the clock regulates *g*_s_ independently from *A*_l_ (Dodd *et al.*, 2014; Dodd *et al.*, 2004). That is, the circadian pattern in leaf carbon assimilation is a function of circadian regulation of leaf biochemistry, and independent of variation in stomatal conductance (Doughty *et al.*, 2006; Dodd *et al.*, 2014; Haydon *et al.*, 2013). Our goal was not to assess the mechanisms driving circadian rhythms in stomata and photosynthesis. However, we note that mechanisms underlying circadian gas exchange regulation are being mostly studied at molecular or cellular scales. Focusing on the mechanisms underlying circadian regulation, at the scales relevant for ecosystem studies, should be at the forefront of our research efforts.

## Concluding remarks

Following conventional wisdom, diurnal variation during the entrainment phases would have been largely attributed to direct environmental effects of PAR, *T*_air_ and VPD on physiological processes (Sellers *et al.*, 1997; Hollinger *et al.*, 1994; Richardson *et al.*, 2007; Jones, 2014; Schwalm *et al.*, 2010). Our experiment using constant environmental conditions as a ‘control’ indicates that up to 79% of the diurnal range in canopy CO_2_ and H_2_O fluxes can be recreated fully independent of environmental change (Fig. 1, Table 1). This diurnal variation under a constant environment showed a period of ~24 h, and can therefore be fully attributed to an circadian controls over leaf photosynthesis (*A*_l_) and stomatal conductance (*g*_s_). Furthermore, we observed how considering circadian rhythms into stomatal models led to improved modeling outputs.

We need additional studies that broadly across phylogenies and functional groups for the expression of circadian regulation in gas exchange. Although current evidence points towards a highly conserved genetic make-up of circadian rhythms plants (Holm *et al.*, 2010), it is still currently unknown under which conditions is circadian regulation of gas exchange expressed (Doughty *et al.*, 2006). Similarly, although our study was performed under radiation levels much higher than those in growth chambers (usually < 200 µmol m^−2^ s^−1^), where the circadian is clock is most often assessed, radiation is still below saturation. We thus need technological improvements that allow achieving saturating radiation loads at ecosystem level (we are unaware of any facility in the world where saturating radiation can be achieved over entire macrocosms or ecosystems while controlling for other environmental drivers).

Our results contribute to the expanding field of plant “memory”, in that the circadian clock regulates gas exchange based upon the conditions of the previous days. Conceptual frameworks on the effects of “memory” on ecological systems often consider the effect of legacies from antecedent environmental stress (Ogle *et al.*, 2015), and potential epigenetic regulations (Crisp *et al.*, 2016). Circadian regulation could acts as an adaptive memory in that a plant's metabolism is adjusted based on the conditions experienced in previous days, and fitness is increased via anticipation (Resco de Dios *et al.*, 2016) and growth regulation (Herrmann *et al.*, 2015; Graf *et al.*, 2010). Our proposed modeling approach expands therefore expands current frameworks on how to incorporate memories from ecological processes into global change models.

## ACKNOWLEDGEMENTS

This study benefited from the CNRS human and technical resources allocated to the ECOTRONS Research Infrastructures as well as from the state allocation 'Investissement d'Avenir’ ANR-11-INBS-0001, ExpeER Transnational Access program, Ramón y Cajal fellowships (RYC-2012-10970 to VRD and RYC-2008-02050 to JPF), the Erasmus Mundus Master Course Mediterranean Forestry and Natural Resources Management (MEDfOR) and internal grants from UWS-HIF to VRD and ZALF to AG. We remain indebted to E. Gerardeau, D. Dessauw, J. Jean, P. Prudent (Aïda CIRAD), J.-J. Drevon, C. Pernot (Eco&Sol INRA), B. Buatois, A. Rocheteau (CEFE CNRS), A. Pra, A. Mokhtar, C.V.M. Barton and the full Ecotron team, in particular C. Escape, for outstanding technical assistance during experiment set-up, plant cultivation or subsequent measurements. Data is freely accessible upon registration from http://www.ecotron.cnrs.fr/index.php/en/component/users/?view=login.

